# The ‘*Candidatus* Phytoplasma mali’ effector protein SAP11_CaPm_ interacts with MdTCP16, a class II CYC/TB1 transcription factor that is highly expressed during phytoplasma infection

**DOI:** 10.1101/2022.07.20.500878

**Authors:** Cecilia Mittelberger, Bettina Hause, Katrin Janik

## Abstract

‘*Candidatus* Phytoplasma mali’, is a bacterial pathogen associated with the so-called apple proliferation disease in *Malus* × *domestica*. The pathogen manipulates its host with a set of effector proteins, among them SAP11_CaPm_, which shares similarity to SAP11_AYWB_ from ‘*Candidatus* Phytoplasma asteris’. SAP11_AYWB_ interacts and destabilizes the class II CIN transcription factors of *Arabidopsis thaliana*, namely AtTCP4 and AtTCP13 as well as the class II CYC/TB1 transcription factor AtTCP18, also known as BRANCHED1 being an important factor for shoot branching. It has been shown that SAP11_CaPm_ interacts with the *Malus* × *domestica* orthologues of AtTCP4 (MdTCP25) and AtTCP13 (MdTCP24), but an interaction with MdTCP16, the orthologue of AtTCP18, has never been proven. The aim of this study was to investigate this potential interaction and close a knowledge gap regarding the function of SAP11_CaPm_. A Yeast two-hybrid test and Bimolecular Fluorescence Complementation *in planta* revealed that SAP11_CaPm_ interacts with MdTCP16. MdTCP16 is known to play a role in the control of the seasonal growth of perennial plants and an increase of *MdTCP16* gene expression has been detected in apple leaves in autumn. In addition to this, *MdTCP16* is highly expressed during phytoplasma infection. Binding of MdTCP16 by SAP11_CaPm_ might lead to the induction of shoot proliferation and early bud break, both of which are characteristic symptoms of apple proliferation disease.

## Introduction

‘*Candidatus* Phytoplasma mali’ (‘*Ca*. P. mali’) is the bacterial pathogen associated with the so-called apple proliferation disease [1]. Apple trees affected by this disease show different growth aberrations like uncontrolled shoot proliferation or enlarged stipules, early bud break and flowering and so-called witches’ brooms. Symptomatic trees produce small, tasteless, and unmarketable fruits, which causes economic losses and threatens apple production [2]. Phytoplasmas are wall-less, phloem-limited bacteria, that need insect vectors for their transmission and use effector proteins to manipulate their host plants [3–6]. ‘*Ca*. P. mali’ is transmitted by certain psyllids [7] and encodes several potential effector proteins that appear to be released into *Malus* plants by a Sec-dependent secretion system [8,9]. So far, the best characterized effector protein is SAP11_CaPm_, which shows similarity to SAP11_AYWB_ from ‘*Candidatus* Phytoplasma asteris’ that is associated with Aster Yellow Witches’ Broom disease in aster [10]. Despite of being the best described effector, SAP11_CaPm_’s function is not fully unraveled yet, and there are still knowledge gaps regarding its interaction partners in *Malus* × *domestica*, the natural host of ‘*Ca*. P. mali’.

The development of witches’ brooms, i.e., uncontrolled branching, is a common symptom not only for apple proliferation, but also for several other phytoplasma diseases. Overexpression of SAP11_AYWB_ and SAP11-like proteins from other phytoplasma species resulted in an increased formation of lateral shoots [11–15]. Interestingly, SAP11_AYWB_, SAP11_CaPm_ and other SAP11-like proteins bind and destabilize different TEOSINTE BRANCHED1/CYCLOIDEA/PROLIFERATING CELL FACTOR 1 and 2 (TCP) transcription factors [12,14,16].

TCP transcription factors are highly conserved in all land plant lineages and contain a basic helix-loop-helix (bHLH) domain, responsible for DNA binding [17]. TCPs are classified according to sequence differences in the bHLH-domain in class I and class II subfamilies [18]. The class II TCPs are further subdivided in CINCINNATA (CIN)-like TCPs and in CYCLOIDEA/TEOSINTE BRANCHED1 (CYC/TB1)-like TCPs [19]. TCPs of both classes play a key role in morphological development of plants, stress adaptions and plant immunity [20] and are thus interesting targets of diverse pathogen effector proteins [21,22].

A total of 52 TCP-domain containing genes were identified in apple [23]. For CIN-like class II TCPs, MdTCP25 (orthologue to AtTCP4) and MdTCP24 (orthologue to AtTCP13), an interaction with SAP11_CaPm_ has been shown [8]. Additionally, a yeast-two-hybrid (Y2H) screen using SAP11_CaPm_ revealed cDNA fragments of MdTCP16, a AtTCP18-like CYC/TB1 class II TCP and of the putative chlorophyllide b reductase NYC1 as putative interaction partners [8]. An interaction between SAP11_CaPm_ and the corresponding full-length gene products of the MdTCP16- and MdNYC1-fragment could not be confirmed at that time. Shortly after, the genome of *Malus* × *domestica* was *de novo* assembled and updated [24]. This sheds new light on the full-length ORFs including *MdTCP* encoding genes and led to some new sequence and reading-frame information regarding the assigned full-length genes of *MdTCP16* and *MdNYC1*.

AtTCP18 is a key regulator of shoot branching [25–27] and is also known as BRANCHED1 (BRC1). BRC1 is used as a common term for all AtTCP18 orthologues in other plant species. Its expression is repressed by cytokinin, gibberellin, phytochrome B and sugar, and is promoted by auxin, an important regulator of apical dominance, strigolactones and a low red to far-red light ratio [28]. BRC1 influences not only the plant architecture, but also the seasonal growth of perennial plants, such as temperate fruit trees by responding to photoperiodic changes [29]. In addition, BRC1 interacts with the FLOWERING LOCUS T protein in Arabidopsis and represses floral transition in axillary meristems [30].

Since several other SAP11-like proteins interact with AtTCP18-like TCPs [11,13,15,31,32] and SAP11_CaPm_ from ‘*Ca*. P. mali’ strain PM19 showed an interaction with AtTCP18 in an Y2H screen [16], the aim of this study was a detailed analysis of the two potential interactions between SAP11_CaPm_ with either MdTCP16 (i.e., the orthologue of AtTCP18) or MdNYC1. Moreover, to gain a better understanding of the role of MdTCP16, MdTCP24 and MdTCP25 in the host plant during phytoplasma-infection, the expression of these *TCPs* was determined by qPCR analysis of healthy and infected *Malus* × *domestica* leaf samples in spring and autumn.

## Materials and Methods

### Yeast-two-hybrid screen

In a previous study cDNA sequences that were partially similar to genes encoding *MdTCP16* and *MdNYC1* were identified as interactors of SAP11_CaPm_ from ‘*Ca*. P. mali’ strain STAA (Accession: KM501063) by a Y2H screen [8]. These cDNA nucleotide sequences were obtained by sequencing the prey vectors in the identified yeast colonies and blasting against the NCBI nt-database [33,34]. The sequence, identified as *MdTCP16*, shared 100% sequence identity with the reference sequence XM_008376500.2 and covers 69.5 % of the coding sequence for protein XP_008374722.1 (S1 Fig). Primers were designed to amplify the coding sequence of XP_008374722.1 (S1 Table). These primers contained overhangs that attached *Sfi*I-restriction sites to the amplicon for subsequent cloning into the pGAD-HA Y2H prey vector.

The sequence identified as *MdNYC1* shared 100% sequence identity to the reference sequence XM_029109831.1 and covers 13.7% of the C-terminal end of XP_028965664.1. Primers were designed to amplify the coding sequence of XP_028965664.1 and of the C-terminal part identified as an interactor of SAP11_CaPm_ in the Y2H screen. Both amplicons were subcloned into pGAD-HA prey vector via their primer-attached *Sfi*I-overhangs.

The Y2H prey vector was co-transformed with pLexA-N-SAP11_CaPm_ bait vector into the *Saccharomyces cerevisiae* strain NMY51 cells [8,35]. Growth was monitored four days after transformation on selective SD plates lacking the amino acids adenine, leucine, histidine and tryptophane.

### Bimolecular fluorescence complementation analysis

Primers, specific for *MdTCP16* and *SAP11_CaPm_* from ‘*Ca*. P. mali’ strain STAA and with *attB*-site overhangs were designed (S1 Table) and fragments were amplified in a total of 50 μL reaction volume using 10 ng template and a final concentration of 1 x iProof HF Buffer, 200 μM dNTPs (50 μM each nucleotide), 0.5 μM of each primer and 0.02 U/μl iProof DNA Polymerase (Bio-Rad, Hercules, CA). The cycling conditions were 30 sec of initial denaturation at 98°C followed by 35 cycles of 98°C for 10 sec, 60 °C for 30 sec and 72°C for 90 sec. The final extension was carried out at 72°C for 5 min. The amplified fragments were analyzed on a 1.2 % agarose-gel and extracted with Kit Montage Gel Extraction columns (Millipore, Bedford, MA). The purified amplicons were used in a BP-reaction for the creation of Gateway-Entry vectors using 1 μL purified amplicon, 100 ng of donor vector (pDONR221-P1P4 or pDONR221-P3P2), 1 μL BP Clonase™ II enzyme (Invitrogen, Carlsbad, CA) following the manufacturer’s protocol. Depending on the N- or C-terminal location of the split-YFP in pBiFC vectors [36], *MdTCP16* and *SAP11_CaPm_* entry-vectors with or without stop-codon were combined in a LR-reaction using Gateway™ using LR Clonase™ II enzyme mix (Invitrogen) and following the manufacturer’s instructions.

*Nicotiana benthamiana* leaf mesophyll protoplasts were isolated for bimolecular fluorescence complementation analysis (BiFC) from leaves of four-week-old plants and protoplasts were transformed with the pBiFC vectors as described in Janik et al. (2017) [37]. Transformed protoplasts were analyzed 16h after transformation using a confocal laser scanning microscope (Zeiss LSM800, Carl Zeiss Microscopy, Oberkochen, Germany). Transformation rate and BiFC rate were determined as a mean value of three independent repetitions of protoplast transformation.

### RNA extraction and cDNA synthesis

Greenhouse samples were collected in May and October 2021. Leaf discs from five non-infected and five infected, one year old apple trees, grown in greenhouse and graft inoculated with ‘*Ca*. P. mali’ infected shoots, were excised, and immediately frozen in liquid nitrogen. Leaf discs were grinded with mortar and pestle under liquid nitrogen. 100 mg of leaf powder was used for RNA extraction following protocol A of the Spectrum™ Plant Total RNA Kit (Sigma-Aldrich, St. Louis, MO). RNA was eluted in 50 μL elution buffer. RNA concentration was measured with a Spectrophotometer (NanoPhotometer® N60, Implen, München, Germany). 1 μg or 2 μg of RNA were reverse transcribed into cDNA by using SuperScript™ IV VILO™ Master Mix with ezDNase enzyme (Invitrogen), following the protocol that includes gDNA digestion with DNase enzyme. Samples that contained all reagents and RNA except the reverse transcriptase (“No-RT controls”) were performed in parallel. Synthesized cDNA was stored at −80°C.

Sampling of naturally infected trees of *Malus* × *domestica* cv. Golden Delicious was done in May and October 2011. RNA extraction and cDNA preparation is described in Supplementary Material and Methods of Janik et al. (2017) [8]. Synthesized cDNA was stored at −80°C.

### DNA extraction

DNA of *Malus* × *domestica* leaf samples was extracted from approx. 50 mg grinded leaf tissue, using the DNeasy Plant Mini Kit (Qiagen, Venlo, The Netherlands), following the manufacturer’s instruction. DNA was eluted in 50 μL elution buffer.

### qPCR analysis

The transcription factors *MdTCP16, MdTCP25, MdTCP24* and the effector protein *SAP11_CaPm_* were amplified with specific qPCR primer pairs (S1 Table) using a SYBR-Green qPCR assay. Reactions were run in a total of 10 μl, using 2 x Universal KAPA SYBR® FAST master mix (KAPABIOSYSTEMS, Wilmington, MA), 20 pmol of forward and reverse primer and 2 μl of 1:50 diluted cDNA samples. Additionally, qPCR master mixes for the reference genes *GAPDH*, *tip41* and *EF1α* were prepared for all samples, using the same reagent and template concentrations, and cycling conditions as mentioned above. All targets were amplified on the same 384well plate in one qPCR run on a CFX384 Touch Real-Time PCR Detection System (Bio-Rad), using three technical replicates per sample. Cycling conditions were as follows: 95°C for 20 sec, followed by 35 cycles with 95°C for 3 sec and 60 °C for 30 sec, melt curve from 65°C to 95°C with an increment of 0.5°C/5 sec.

For determining the qPCR efficiency of every primer-combination, a four-point dilution series (1:10, 1:50, 1:100, 1:200) of a cDNA sample mixture was analyzed in each qPCR run.

Phytoplasma quantity was determined by the detection of the phytoplasma specific *16S* gene together with the *Malus* specific single copy gene *ACO* as described by Baric et al. (2011) [38]. In brief, 2 μl of template DNA were analyzed in a total multiplex reaction volume of 20 μl, using 2x iQ™ supermix master mix (Bio-Rad), 18 pmol of each qAP-16S forward and reverse primer, 4 pmol qAP-16S probe, 4 pmol of each qMD-ACO forward and reverse primer and 4 pmol of qMD-ACO probe. The ‘*Ca*. P. mali’ specific qAP-16S probe was 5’-labeled with the reporter dye FAM, while the *Malus* specific qMD-ACO probe was 5’-labeled with VIC. Cycling conditions were as follows 95°C for 3 min followed by 35 cycles of 95°C for 15 sec and 60 °C for 60 sec. Reactions were run in triplicates on a CFX96 Touch Real-Time PCR Detection System (Bio-Rad). For determining the qPCR efficiency, a five-point dilution series (undiluted, 1:10, 1:50, 1:100, 1:200) of a sample mixture of DNA from infected *Malus* × *domestica* roots was analyzed in each qPCR run.

### Data analysis

Normalized expression was calculated according to Taylor et al. (2019) [39] considering qPCR efficiency (E) of each run, following the formula 1+E^ΔCq^ for relative quantity. Statistical analysis was performed with GraphPad Prism 7.05 (GraphPad Software Inc., La Jolla, CA).

## Results

### Yeast-two-hybrid and bimolecular fluorescence complementation analyses

The Y2H test using SAP11_CaPm_ from ‘*Ca*. P. mali’ strain STAA fused to the GAL4-binding domain as bait revealed an interaction between MdTCP16 and SAP11_CaPm_. An interaction between SAP11_CaPm_ and MdNYC1 (neither the fragment nor the full-length protein) was not detectable (Fig 1).

**Fig 1.**
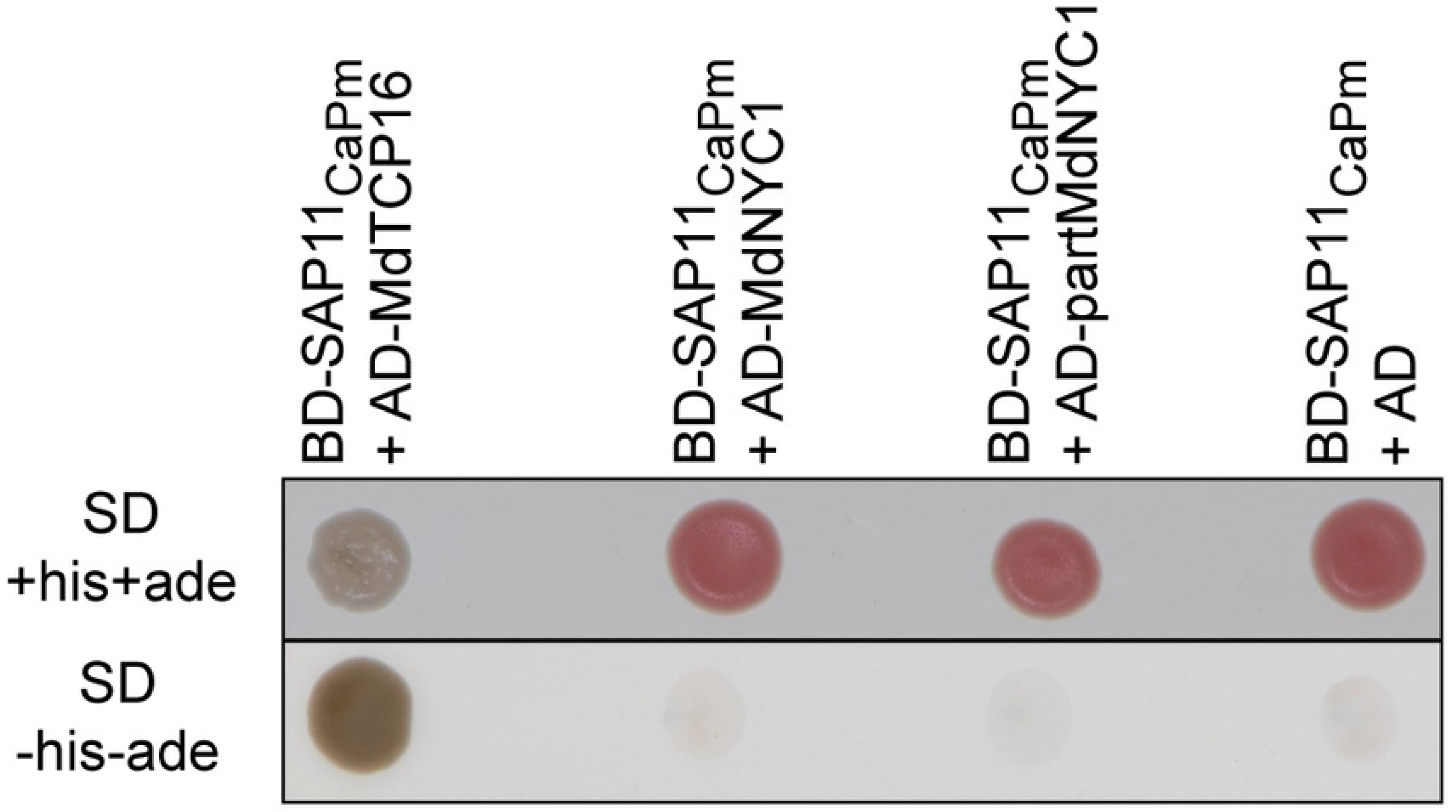
SAP11_CaPm_ interacts with MdTCP16 but not with MdNYC1 in yeast. Interaction between the bait SAP11_CaPm_ fused to a DNA binding domain (BD) and the prey protein from *Malus* × *domestica* fused to an activation domain (AD) is indicated by growth of the *Saccharomyces cerevisiae* reporter strain NMY51 on SD minimal medium lacking the amino acids histidine (his) and adenine (ade). The white color of BD-SAP11_CaPm_ + AD-MdTCP16 grown on full medium is an additional indication for a strong interaction, since the ADE2 reporter gene is activated upon interaction, while in absence of a protein-protein interaction and thus no ADE2 activation, a red colored intermediate accumulates in the adenine metabolic pathway.

For further confirmation of the interaction between SAP11_CaPm_ and MdTCP16, the sequences of SAP11_CaPm_ and MdTCP16 were subcloned into different pBiFC-2in1 vectors [36] by Gateway-cloning [40]. The pBiFC-2in1 vectors were transformed into *N. benthamiana* mesophyll protoplasts [37] and interaction was analyzed by confocal microscopy 16 h after transformation. Up to 92.5% of the transformed protoplasts showed a YFP signal, resulting from the interaction between SAP11_CaPm_ and MdTCP16 (Fig 2). The YFP-signal was localized to the cell nucleus and occurred additionally in the cytoplasm. The co-expression of SAP11_CaPm_ and MdNYC1 fused to both YFP-halves served as a negative control and did not reveal YFP fluorescence after transformation of protoplasts (Fig 2).

**Fig 2.**
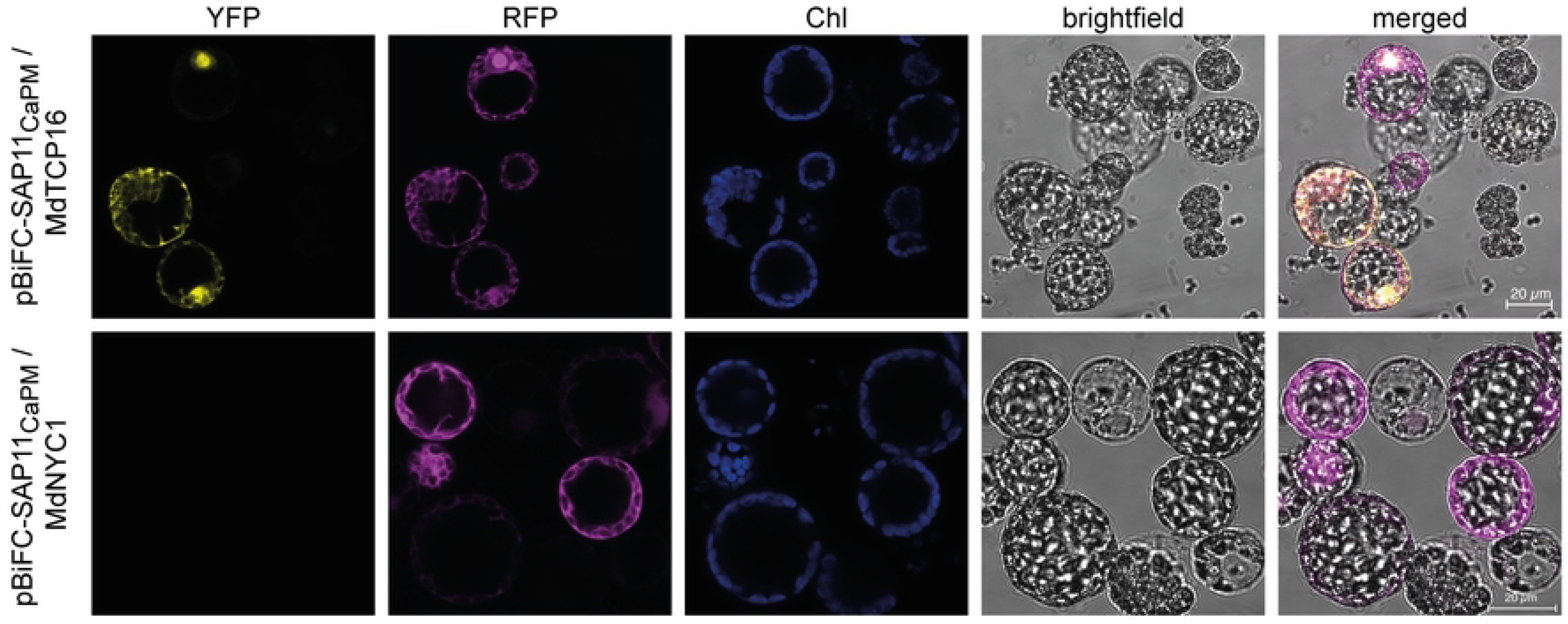
SAP11_CaPm_ interacts with MdTCP16 *in planta*. *Nicotiana benthamiana* mesophyll protoplasts were co-transformed with a BiFC expression vector encoding SAP11_CaPm_ and MdTCP16. SAP11_CaPm_ and MdTCP16 interact in the nucleus and in the cytoplasm as indicated by the occurrence of a YFP signal in these two cellular compartments. The co-expression of SAP11_CaPm_ and MdNYC1 served as a negative control. The RFP signal (depicted in magenta) indicates a successful transformation of protoplasts with the BiFC vector. Chl (depicted in blue) shows the autofluorescence of chlorophyll within the chloroplasts. Microscopic analysis was performed with a Zeiss LSM 800 confocal microscope. Bars represent 20 μm for all micrographs.

Taken together, these results show that SAP11_CaPM_ interacts with MdTCP16 in addition to the previously shown interaction with MdTCP24 and MdTCP25 [8].

### Expression of TCPs in *Malus × domestica* during infection

It was unclear how an infection with ‘*Ca*. P. mali’ affects the expression of the three TCP-encoding genes being the targets of its effector SAP11_CaPm_. Thus, to analyze the expression of *MdTCP16*, *MdTCP24* and *MdTCP25* during infection in *Malus* × *domestica* host plant, qPCR assays were used to determine expression levels of the respective TCPs and the effector protein SAP11_CaPm_.

*MdTCP24* was stably expressed throughout the season and its transcript levels did not change upon infection, regardless of whether the trees were grown in the field or in the greenhouse (Figs 3A and 3B). Also, *MdTCP25* in leaves from greenhouse plants was stably expressed throughout the season, but in naturally infected samples the expression was significantly higher in samples taken in spring than in those taken later in the season, for both healthy and infected trees. Furthermore, regarding field-grown trees in spring, *MdTCP25* expression is significantly lower in naturally infected samples compared to healthy samples (Figs 3A and 3B). The expression of *MdTCP16* was lower than those of *MdTCP24* and *MdTCP25. MdTCP16* transcript levels were low in both infected and healthy samples from spring and higher in samples from autumn. Furthermore, it was tendentially, but not significantly higher in infected autumn samples than in healthy autumn samples (Figs 3A and 3B). *SAP11_CaPm_* expression could not be detected in naturally infected spring samples, but it was detectable in the leaves from greenhouse samples throughout the season (Fig 3C). This discrepancy might be due to the season-dependent phytoplasma colonization pattern of apple trees [38]. SAP11_CaPm_ expression in leaves tends to be increased when phytoplasma level in the same tissue is high (Fig 3D).

**Fig 3.**
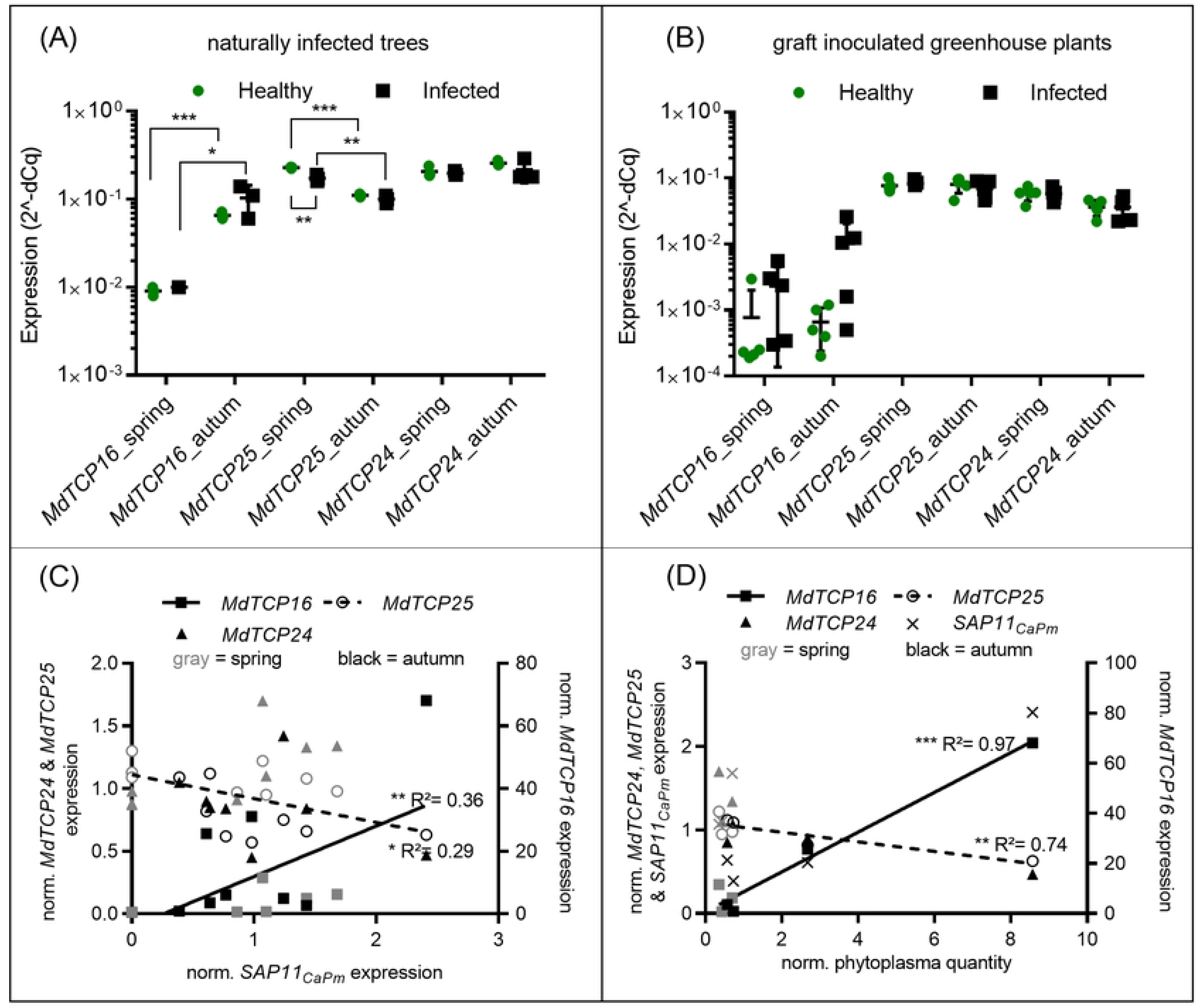
*MdTCP16* expression increases from spring to autumn, is slightly higher in infected samples and strongly correlates to phytoplasma levels in leaves from infected *Malus × domestica*. MdTCP expression in spring and autumn of non-infected (green) and naturally infected (black) apple leaves (A) or graft inoculated and grown in greenhouse plants (B). Correlation of *MdTCP* and *SAP11_CaPm_* expression (C) and phytoplasma levels (D) in infected *Malus* × *domestica* leaf-samples from spring (grey) and autumn (black). Statistical analysis was performed with multiple t-test. Statistical significance was determined using the Holm-Sidak method, with alpha = 0.05 and linear regression analysis, using GraphPad Prism 7.05 (GraphPad Software Inc.). Significant differences between groups are indicated with asterisks (* P≤0.05, ** P≤0.01, *** P≤0.001).

The expression of *MdTCP16* correlates moderately positively (R^2^=0.36) and strongly positively (R^2^=0.97) with the expression of *SAP11_CaPm_* and the phytoplasma quantity in leaf samples, respectively (Figs 3C and 3D, Table 1). This is opposite to *MdTCP25*, whose expression correlates weakly negatively (R^2^=0.29) and negatively (R^2^=0.74) with the expression of *SAP11_CaPm_* and the phytoplasma quantity in leaf samples, respectively (Figs 3C and 3D, Table 1). *MdTCP24*, in contrast, does not correlate with the expression of *SAP11_CaPm_* nor with the phytoplasma quantity (Figs 3C and 3D).

**Table 1.**
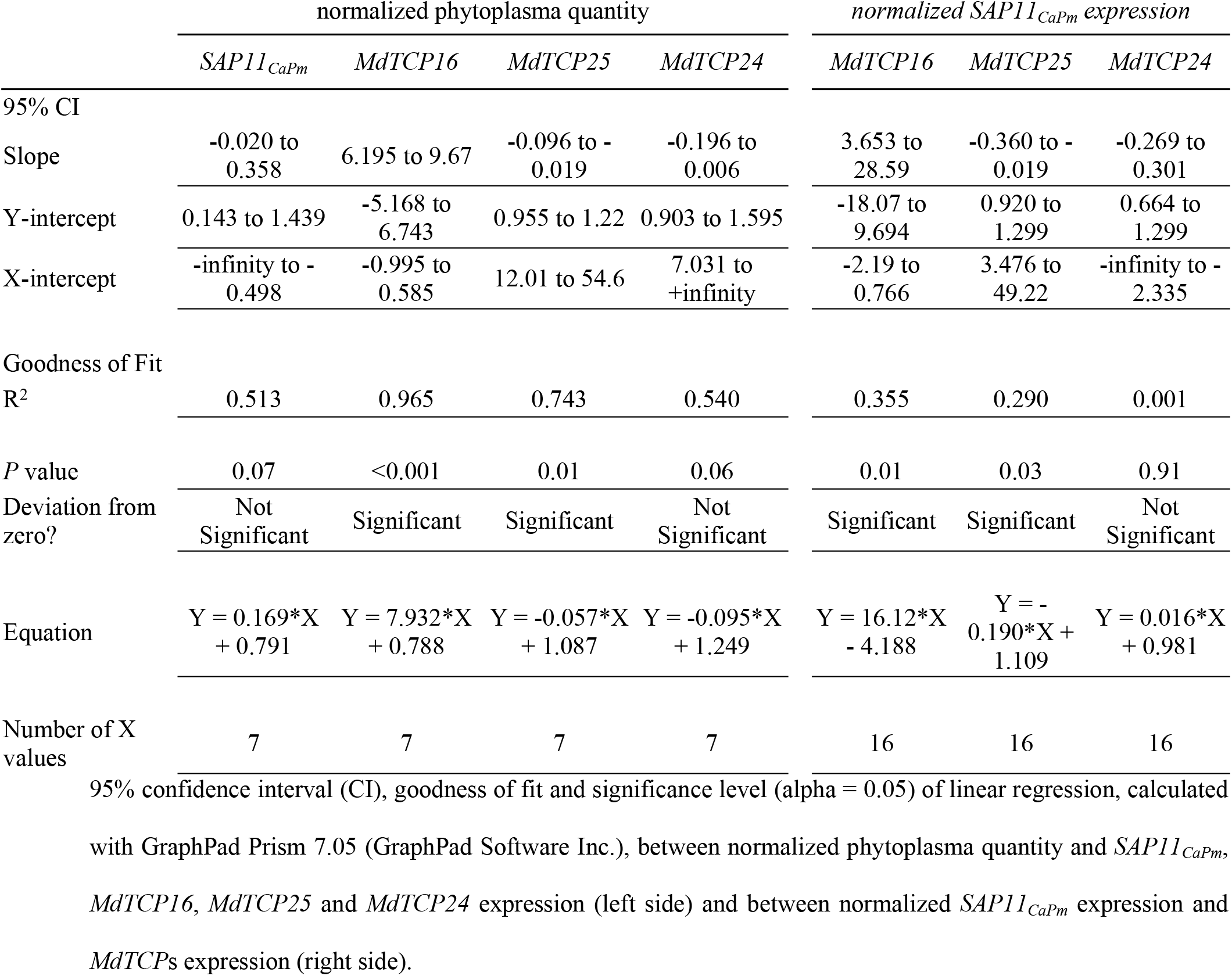
Linear regression analysis of TCP expression depending on phytoplasma quantity or *SAP11_CaPm_* expression.

Concluding from these analyses, only the expression of *MdTCP16* in leaves is positively correlated with infection, thereby most highly expressed late in autumn.

## Discussion

It is known that SAP11_AYWB_ from AYWB-phytoplasma interacts and destabilizes AtTCP18 in *Arabidopsis thaliana* [12], but it was so far unknown if SAP11_CaPm_ from ‘*Ca*. P. mali’ can interact with MdTCP16, the orthologous of AtTCP18 in *Malus* × *domestica*, the actual host plant of this phytoplasma species. It has been shown previously that SAP11_CaPm_ from strain PM19, that shares 99.2% sequence identity to SAP11_CaPm_ from strain STAA used in this study, interacts with AtTCP18 in a Y2H test [16]. However, to understand the potential effects of the effector protein SAP11_CaPm_, it is important to unravel the targets not only in model plants but in the natural host plant and to understand the induced changes in TCP expression in the native pathosystem.

Our data show unequivocally that SAP11_CaPm_ interacts with MdTCP16, both yeast and in planta, whereas another candidate (MdNYC1) does not interact (Fig 1, 2). SAP11_CaPm_ also binds to MdTCP25 and MdTCP24 from *Malus* × *domestica* and degrades different AtTCPs at the protein level, among them the orthologue of MdTCP25 i.e., AtTCP4 [16], but it is not known, whether binding of SAP11_CaPm_ affects the stability of MdTCP16, since it was not able to destabilize the *Arabidopsis thaliana* orthologue AtTCP18 [31]. Interestingly, our study indicates that *MdTCP16* expression is induced during phytoplasma infection and might then be counter-regulated by SAP11_CaPm_ at the protein level. A degradation of MdTCP16 induced by SAP11_CaPm_ at the protein level could induce *MdTCP16* expression on the transcriptional level via a gene-product mediated feedback loop regulation [41]. The MdTCP16 protein might act like a repressor and negatively affects expression of the *MdTCP16* gene, meaning that skimming the proteinaceous gene product leads to an increase of its gene expression. That might explain why *MdTCP16* transcriptional expression levels are tendentially increased in infected plants (Fig 3A and 3B) which has been shown also for ‘*Ca*. P. ziziphi’ infected *Ziziphus jujube*, another phytoplasma pathosystem where the *ZjTCP7* that encodes a MdTCP16 orthologue of *Z. jujube*, is upregulated in leaves [42]. It is worth noting that – in contrast to *MdTCP16* – *MdTCP25* expression is adversely i.e., negatively and *MdTCP24* expression not at all correlated with SAP11_CaPm_ expression during infection. This implies that other regulatory mechanisms might be involved in the regulation of *MdTCP25* and *MdTCP24* expression.

Whatever pathway is affected during *MdTCP16* expression, it seems reasonable to speculate that it is important for the phytoplasma to attack MdTCP16 in its plant host.

The increased *MdTCP16* expression in autumn samples might be an indication that this TCP is involved in the seasonal control of branching. *MdTCP16* is mainly expressed in axillary and flower buds and only weakly in leaves, stems and shoot tips of apple trees [43]. Short photoperiods lead to the expression of *BRC1* and together with low temperature a complex network controls bud dormancy [44]. Interestingly, early bud break is a symptom of Phytoplasma-infected apple trees. Therefore, it can be assumed that the effect of ‘*Ca*. P. mali’ infection on MdTCP16 (a member of the *BRC1* orthologue group) function might be even more pronounced in axillary buds and the binding of SAP11_CaPm_ to MdTCP16 might decrease the quantity of this TCP in axillary buds and thus trigger the earlier bud break in spring.

‘*Ca*. P. mali’ infection leads to increased soluble sugar content in phloem sap [45] and in leaves [46]. Sugar promotes the bud outgrowth by acting as a repression-signal for *BRC1* expression [25,47]. This is neither in line with the increased *MdTCP16* expression that we find in phytoplasma infected apple leaves, nor with what has been described for *ZjTCP7* expression in phytoplasma infected Chinese jujube [42]. Thus, other factors might outcompete the repressing effect of increased sugar levels on *MdTCP16* expression, such as an increased auxin level in leaves of infected apple trees [48]. Auxin induces the *MdTCP16* expression but blocks the axillary bud outgrowth in healthy plants [49,50].

Taken together, BRC1 and its homologues seem to be important molecular targets of SAP11-like effector proteins from different phytoplasma species. The results of this study prove the interaction of SAP11_CaPm_ and the MdTCP16 transcription factor. SAP11-like proteins might be crucial for the successful phytoplasma colonization of the canopy in spring by downshifting BRC1. In the context of the current knowledge, it can be assumed that this downregulation is involved in the formation of lateral shoot outgrowth and early bud break, which are typical symptoms of ‘*Ca*. P. mali’ infection, and eponymous to the diseases name “apple proliferation” [31]. However, the factors involved in *BRC1/MdTCP16/AtTCP18* gene regulation are not easy to detangle, since they seem to be species- and tissue-dependent and regulated in a complex manner. By expressing SAP11-like proteins, phytoplasma target different members of the plant TCP family which serve as molecular hubs, to manipulate their plant hosts in a very sophisticated -but not yet fully understood-manner.

## Acknowledgments

We would like to thank Christine Kerschbamer for lab assistance and Mirko Moser (Fondazione Edmund Mach, San Michele All’Adige, Italy) for discussing the data. Furthermore, we would like to thank Cameron Cullinan for English proofreading.

## Supporting information

**S1 Fig. Reference Sequence of XM_008376500.2.** The sequence includes the CDS for XP_008374722.1 (*MdTCP16*) with TCP domain and the identified part in the Y2H screen. Graph was generated with Geneious R11.1.5.

**S1 Table. Primers used in this study.** Lowercase letters indicate bases for *Gateway-attB* site overhangs or *Sfi*I restriction site overhangs.

